# A possible origin of the inverted vertebrate retina revealed by physical modeling

**DOI:** 10.1101/2022.10.04.510789

**Authors:** Jan M.M. Oomens

## Abstract

There is no established explanation for the evolution of the inverted retina in the vertebrate camera-type eye. This report shows that the inverted retina may have evolved in the light-sensitive organ of the predecessor of the vertebrates. A patch of light-sensitive epithelium with light-facing photoreceptors developed through evagination into a transparent spherical organ covered in photoreceptors. Photons can activate a photoreceptor either directly or indirectly through the transparent interior of the organ. Models for photoreceptor distribution and spatial resolution show how the inverted retina may have evolved with minimal morphological changes, assuming that natural selection driven optimization of the maximum detectable spatial resolution played a major role.

## Introduction

The eye is a highly complex organ comprising many components and fully understanding the direct evolutionary process that produced it has long eluded scientists. The evolutionary process has to start with light detection, marking the retina as the most ancient part of the eye, whereby two main types occur in the animal kingdom.

The first, well-studied type of retina features light-facing photoreceptors (so-called verted retina) and is found in the eye of flatworms and molluscs, in contrast to the second type of retina in vertebrate eyes, in which the photoreceptor cells are facing away from the light (so-called inverted retina). This appears to be an illogical development, raising the question of how this design may have emerged.

The evolutionary development of the mollusc eye has been sufficiently unraveled (Land & Nilsson, 2012). Nilsson (2009) distinguishes four phases of eye evolution and has the evolutionary process start from a flat area of skin with light-sensitive epithelial cells that perceive local light levels. Through cup-shaped invagination, this area develops directivity to light and its perceptible spatial resolution increases, enabling organisms to optically perceive their surroundings. The emergence of eye appendages such as the cornea, the lens and the pupil in the iris and the increase in the number of receptors in the retina lead to an ever better spatial resolution and thus a more detailed picture of one’s surroundings, improving the organism’s chances of survival (Land & Nilsson, 2012).

For the vertebrate camera-type eye, it remains unclear how the inverted retina might have evolved. Carreras (2018) concluded from his comparative study of the embryogenesis of the eye – involving morphogenesis and Pax6 genetic activity – that the inverted retina was already available in the original ancestor of the vertebrates before neurulation occurred. However, it is still unclear how the inverted retina might have gradually evolved.

There is ample reason to explore a hypothetical evolutionary pathway that could explain the inverted orientation of photoreceptors in the retina of the predecessor of the vertebrate eye. Photoreceptor development and color detection will not be treated here.

This hypothetical evolutionary pathway should eventually lead to an organ enabling accurate visual detection and orientation.

## Method

In the present study, three mathematical models are developed: one model to describe the local photoreceptor density in the retina during the evagination process, a model to calculate the maximum blur spot size on the retina, and a model to estimate the maximum detectable spatial resolution for a spherical light-sensitive organ.

To improve our understanding of the local photoreceptor density, geometry is used to describe how photoreceptor distribution changes during morphological development from the flat light-sensitive patch to the spherical successor. Once the organ has reached its spherical shape, the second model is developed to determine the optical quality and size of the optical blur spot on the retina.

To assess the corresponding detectable spatial frequency in a spherical light-sensitive organ, the theory of Nilsson and Pelger (1994) is applied in model 3. The three models together provide good indicators of how the inverted retina may have originated in the vertebrate eye.

The first phase of eye development is a flat light-sensitive layer with photoreceptor cells oriented towards the light. This layer is covered by a transparent epithelial layer, with basal membranes separating the layers.

There are two opposite ways to easily achieve directional light detection, namely by invagination and evagination of a flat light-sensitive patch. Figure 1 provides a schematic overview of both process.

**Figure 1.**
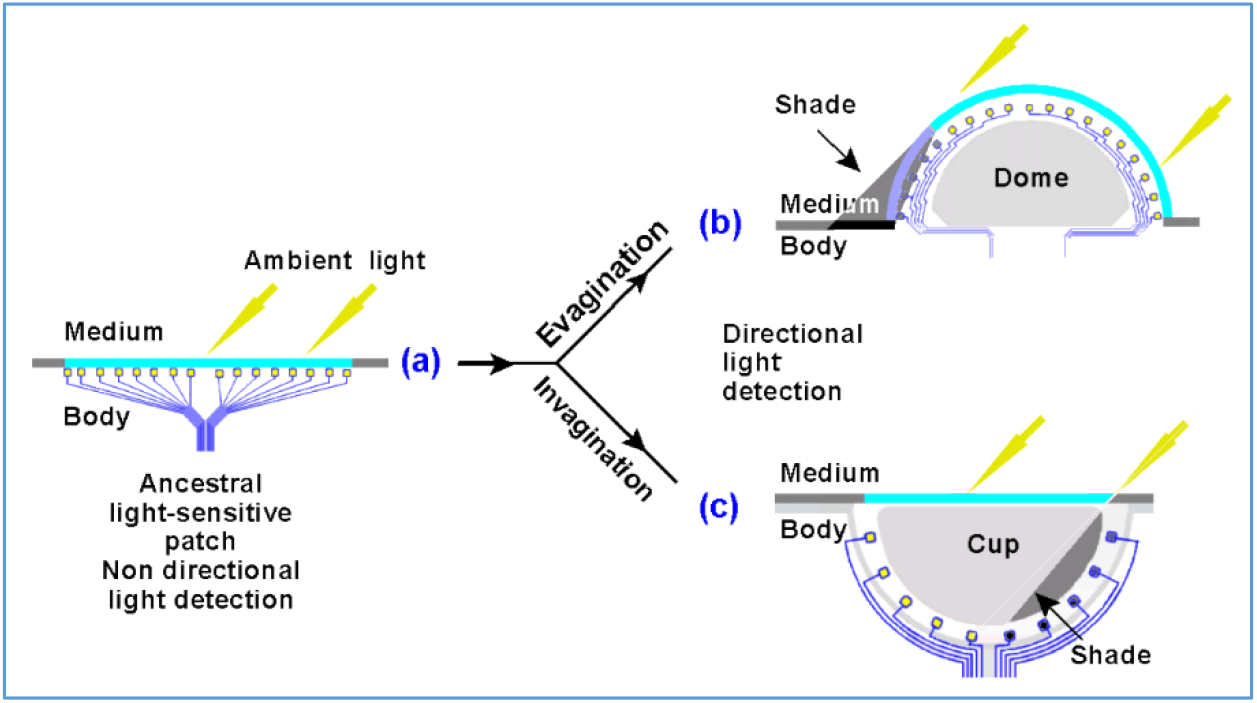
Dome and cup configuration create directional light detection. The light-sensitive patch **(a)** features primitive photoreceptor cells - directly connected to nerves - distributed over its surface. It has two ways to create an organ receptive to the direction of light. By evagination **(b)** forming a dome and invagination **(c)** forming a cup. Both configurations create a shade through refraction and shielding The positions of the optic nerve pathways (blue lines) are considered to remain close to the light-sensitive surface. The nerve pathways are omitted in further figures.

The invagination scenario (c) in Figure 1 is well described by Nilsson and Pelger (1994), Dawkins (1997), and Land and Nilsson (2012) in respect to the evolutionary development of the mollusc eye.

The opposite convex curving of a patch of photoreceptors is mentioned by Dawkins (1997) in relation to the eventual development of compound insects eyes where photoreceptor cells are radially recessed in individual ommatidia, allowing each ommatidium to perceive a portion of the outside world at a small solid angle and achieve spatial resolution.

The evagination scenario (b) in Figure 1 in which a flat light-sensitive patch evaginates towards a spherical structure is not yet properly researched and therefore explored in this study to, eventually, find an explanation for the evolution of the inverted retina of the vertebrate eye.

This research keeps the photoreceptors at the surface of the bulging structure. As the light-sensitive area becomes more convex, it gradually covers an increasingly large field of view and allows the sensory nerve system to develop sensitivity for the direction of light. It finally develops into a light-sensitive sphere.

Figure 2 shows the phases of the hypothetical evolutionary development of the light-sensitive sphere where each phase introduces a new trait that improves the animal’s chances of surviving. The stalk extends the light-sensitive organ away from the body and magnifies the field of perception (Weihs & Moser, 1981), while enabling ground-dwelling creatures to perceive prey and threats from above.

**Figure 2.**
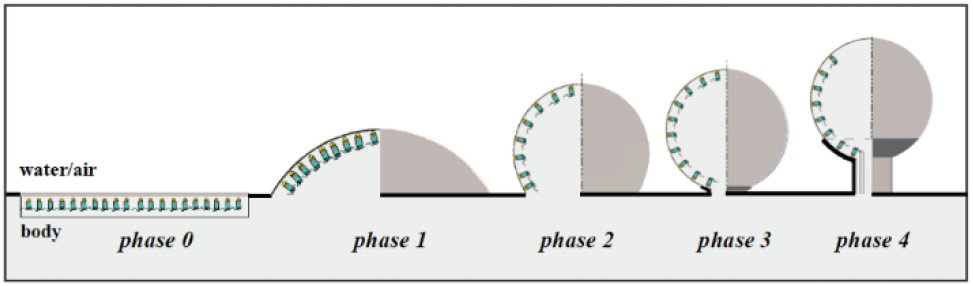
Diagram showing the transformation of a flat light-sensitive patch into a spherical shaped organ on a stalk. The photoreceptors in the patch are oriented to the light. The photoreceptor layer is supported by a transparent vitreous body when it protrudes into the medium, enlarging field of vision. In the successive phases, the vitreous body gains refraction properties of a lensball. The phases are as follows: 0. Round patch of light-sensitive skin, non-directional light detection. 1. Dome-shaped photoreceptor area, directional light detection. 2. Lensball formed due to bulbous necking, posterior retina development. 3. Inverted posterior retina activation and start of stalk development. 4. Forming of an outer light-tight layer, development of a primitive light-sensitive organ with low spatial vision.

In phase 2, the constriction and formation of a primitive light-sensitive spherical organ begins. There is no spatial resolution or imaging at this phase, and only intensity and light direction are observed.

In phase 3, the transparent light-sensitive organ can be understood to be a fully developed lensball sparely covered by photoreceptors. For direct light, the sphere has a convex shaped retina and for incident light a concave retina.

The remarkable feature of the spherical, transparent, light-sensitive organ in phase 4 is that it behaves as a verted retina for the light directly falling on it, as well as an inverted retina for incident light. Two symmetrical light-sensitive organs potentially enable animals to perceive movement (Land & Nilsson, 2012).

The transparent lensball is subject to the laws of optics, which means that photons entering the sphere anteriorly can activate the photoreceptors in the opposite posterior retinal region.

It is unlikely that photoreceptor coverage is entirely uniform across phases, as photoreceptor density will become dense at the base of the sphere between phases 2 and 3 in particular. To rebalance photoreceptor distribution, significant photoreceptor migration in the support matrix will be necessary. It is therefore assumed that photoreceptors retain their position within the support structure of epithelial cells during the evagination process and that shape change is achieved by either the intermediate epithelial cells changing their shape or adapting the support structure to the spherical shape via cell division or cell annihilation. The photoreceptor density distribution on the spherical organ will be very different from the initial density distribution on the light-sensitive patch.

### The photoreceptor density model

The evaginating patch is conceived as a circular membrane that rotationally symmetrically deforms into a light-sensitive spherical surface that protrudes into the medium through which it receives photons. The convex retina is supported by a transparent vitreous structure that - due to its spherical shape - has the refractive properties of a lens.

In photoreceptor density modeling, the light-sensitive patch features photoreceptor cells evenly distributed over its surface. To calculate the photoreceptor density distribution during morphogenesis, the initially flat photoreceptor surface is conceived as a plastically deformable membrane (Figure 3).

**Figure 3.**
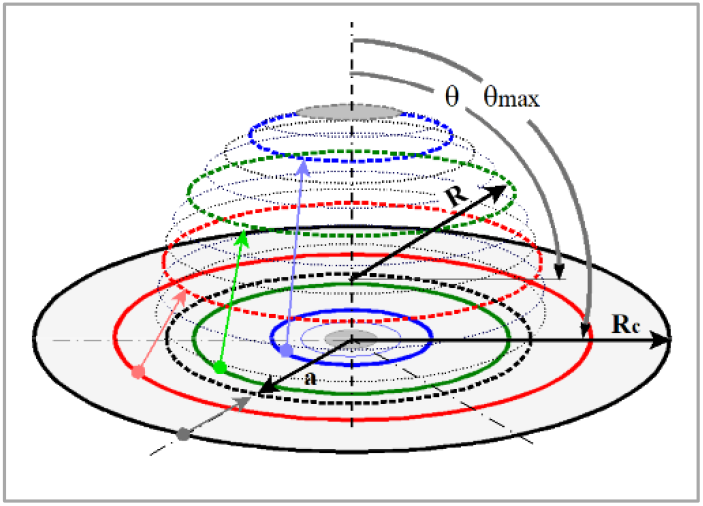
Schematic representation of the parameters involved in the transition of the flat circular light-sensitive patch to a hollow spherical cap. The rim of the patch forms the edge of the spherical cap. The patch (solid lines) has a fixed radius **R_c_**. The spherical cap (dotted lines) has a radius **R**, a cap base diameter of **2a** and a spanned angle **θ_max_** defined from the apex of the cap to its base. The polar angle **θ = 90°** marks the equatorial plane of the spherical cap. The colored rings schematically represent the initial and final positions of photoreceptors on that ring. Depending on the values of the three parameters, local tissue stretching and/or tissue shrinkage occurs. If the arc length over the dome **R·θ_max_ > R_c_**, the distance between photoreceptors on a meridian will increase proportionally and so in the apex. Relative to the apex, the distances between photoreceptors on the parallels will continue to decrease from the apex to the base of the spherical cap. Together, these two effects alter the local photoreceptor density.

Figure 4, Box 1 states the derivation and equation for calculating the local normalized photoreceptor density on the surface of the developing spherical cap. The photoreceptor density distribution is a function of the circle radius *R_c_*, sphere radius *R*, angular position θ and the spanned angle *θ_max_* and is given in equation 1 (Figure 4. Box 1).

**Figure 4.**
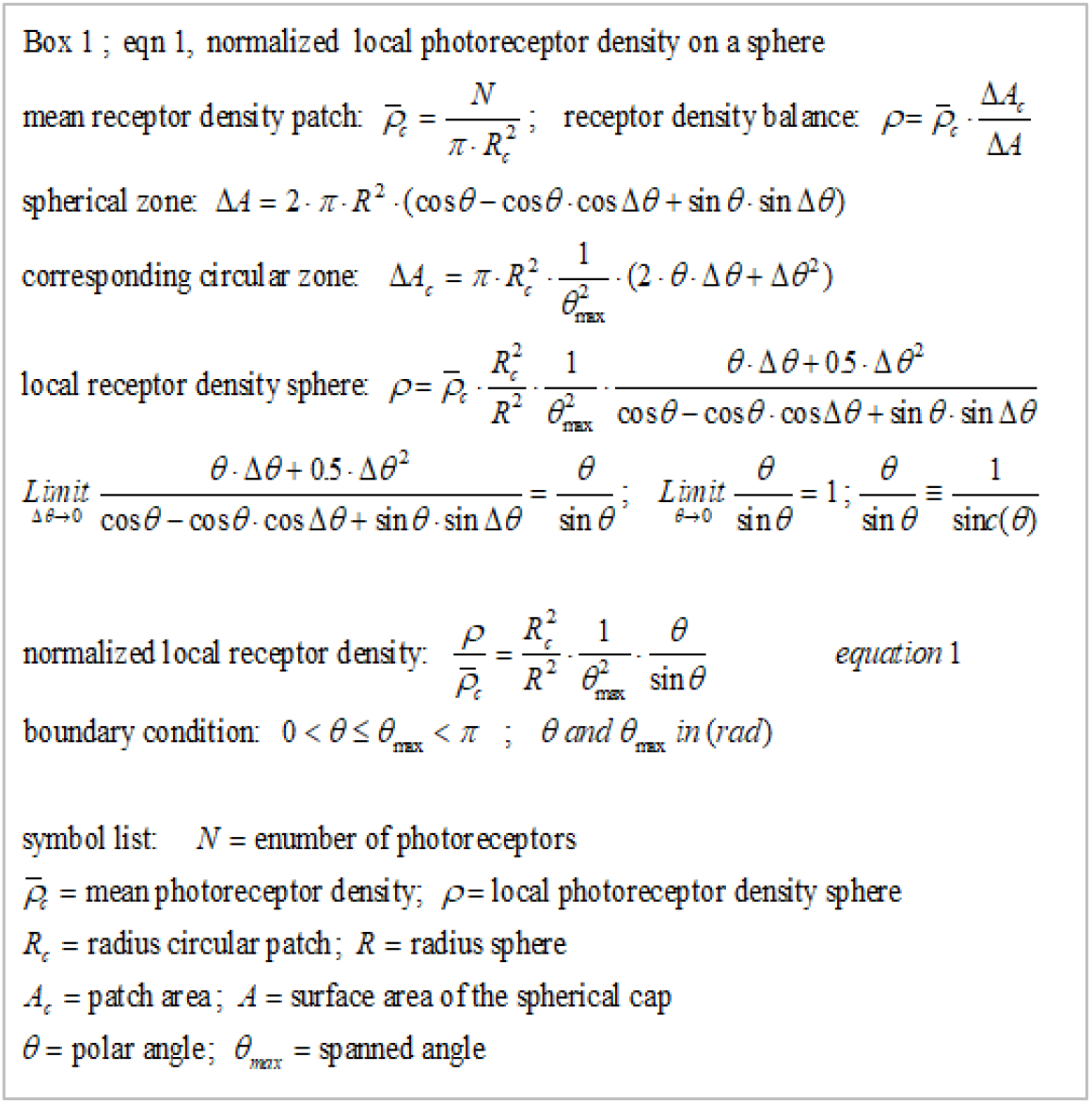
Box 1, Normalized local photoreceptor density on a sphere. **R_c_, R** and **θmax** determine the final state of the evagination process and are therefore boundary conditions for equation 1 within which the local photoreceptor density can be calculated as a function of **θ** (radians). The circle radius **R_c_** and 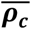. are assumed to be constant and used as fixed reference values so that the local photoreceptor density on the lensball can be presented in a normalized manner. There is a relation between **R_c_** and **R** that depends on the evagination scenarios that will be described in Figure 5.

A scenario in which the radius of the light-sensitive patch remains the same as the radius of the developing spherical cup is used by Nilsson and Pelger (1994). This is a reason to analyze the photoreceptor density on a developing spherical cap for the constant radii scenario.

Equation 1 shows that a smaller sphere radius can provide a greater photoreceptor density and thus better spatial resolution. Maintaining the surface of the circular light-sensitive area and the surface of the spherical cap equal during the bulging process results in a radius *R* becoming increasingly smaller as the spanned angle *θmax* of the spherical cap increases. The constant radius scenario and constant surface area scenario are shown in Figure 5.

**Figure 5.**
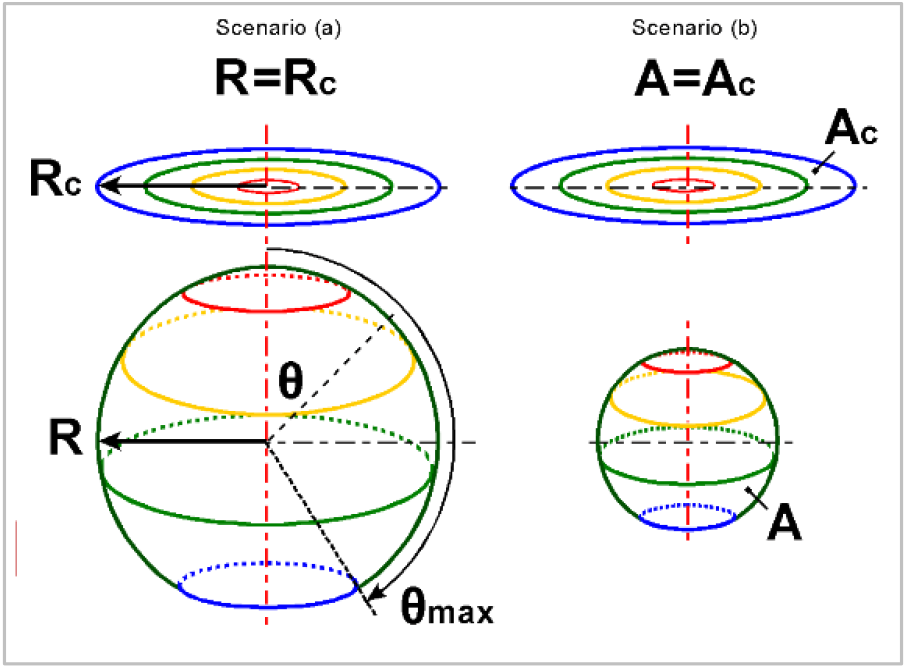
Scale diagram of two evagination scenarios. The initial flat photoreceptor surface and the photoreceptor surface after the development of the spherical photoreceptor organ. The spanned angle **θ_max_** is **170°**. Scenario (a) is the constant radius scenario where **R = R_c_** and **A ≈ 3.9 A_c_.** Scenario (b) is the constant surface area scenario where **A = A_c_** and **R ≈ ½ R_c_.** In scenario (a), the volume of the sphere is eight times the volume in scenario (b).

The corresponding local normalized photoreceptor density equations can be found in *Box 2^1^ (Figure 6*).

**Figure 6.**
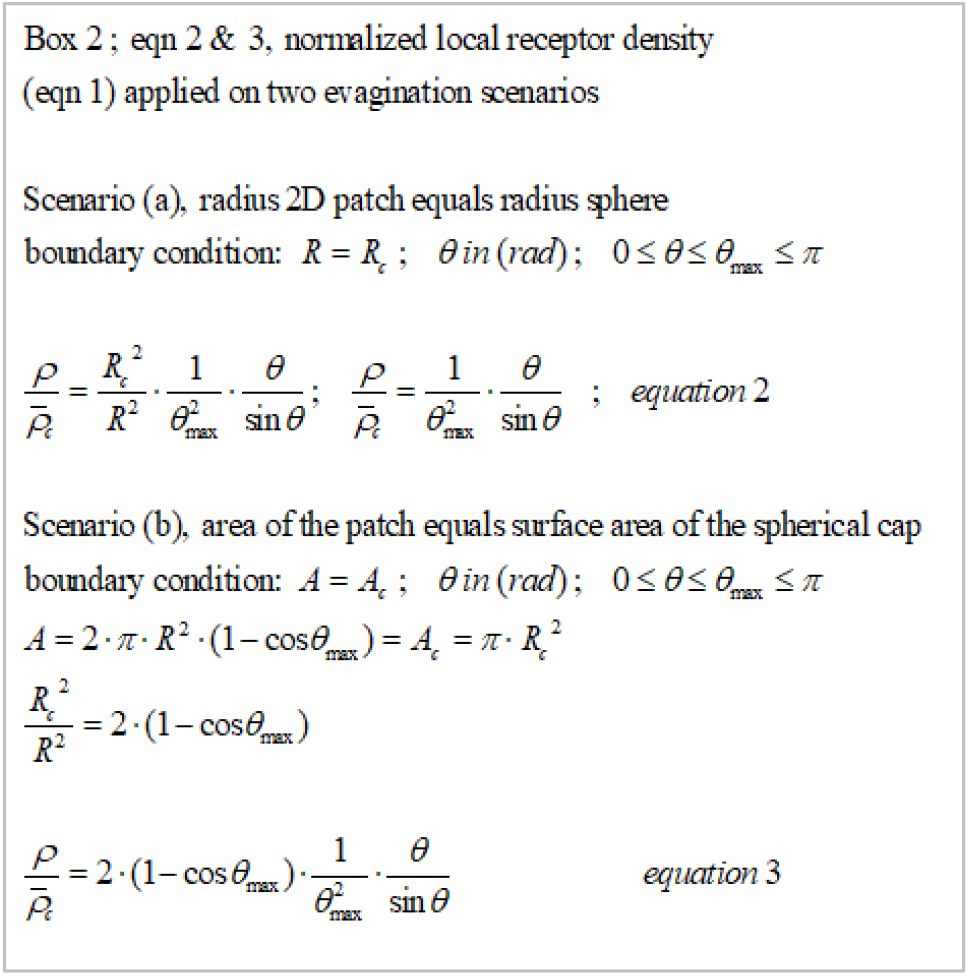
Box 2. Normalized local photoreceptor density equations. Equation 1 applied to the constant radius scenario **R = R_c_** produces equation 2 and to the constant surface area scenario **A = A_c_** equation 3.

The initial photoreceptor density on the light-sensitive patch can be considered to be evenly distributed, whereby it can be attributed a normalized value of 1. With increasing evagination, going along with an increase of the spanned angle θ_*max*_, a dramatic redistribution of the photoreceptor density occurs with a low photoreceptor density in the anterior region and a high density at the base of the lensball surface.

Figure 7 uses equations 2 and 3 to report the local normalized photoreceptor density at a critical spanned angle θ_*max*_ that just allows for the passage of optic nerves and veins through the retina to the connecting stalk, which is necessary for the nutrition of the retina and its communication with a coordinating “brain” system.

**Figure 7.**
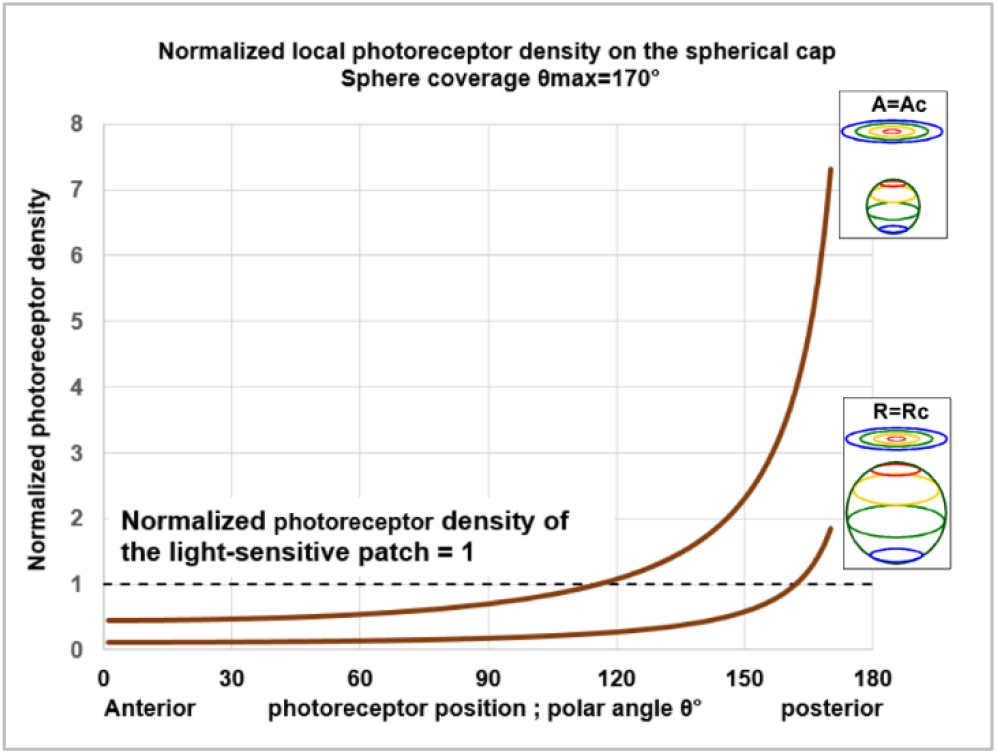
Normalized photoreceptor density profile for two evagination scenarios. A spanned angle **θ_max_** of **170°** is assumed to allow for optic nerves and veins to pass through the retina to the stalk.

In the case of the evagination scenario R=R_c_, the values range from ~0.11 at the anterior part of the sphere untill 1.8 at its base, and for A=A_c_, the values range between ~0.45 and 7.3, respectively. In either case a difference of a factor ~16 in photoreceptor density is observed between the top and base of the spherical cap surface. This phenomenon is further illustrated in a computer-simulated impression of the position of photoreceptors before and after evagination (Figure 8).

**Figure 8.**
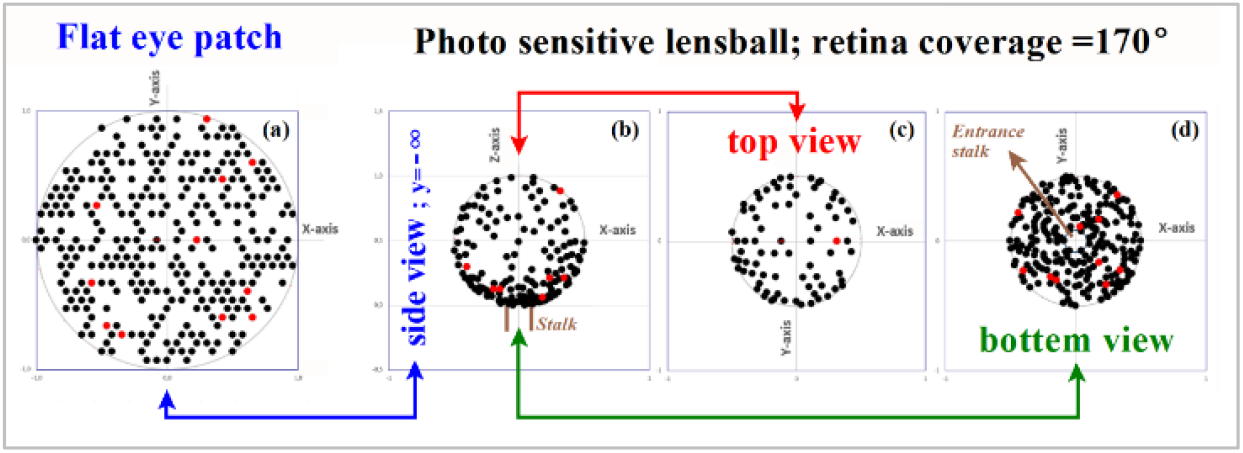
Computer simulation of how the 2D flat receptor cell configuration distributes spatially across the lensball. The flat light-sensitive patch shown in image (a) has 300 photoreceptors randomly placed in a hexagon lattice with 600 positions. Images (b), (c) and (d) give an impression of the individual photoreceptor positions in the evagination scenario **A=A_c_**. The red dots are markers to show how individual photoreceptors from the light-sensitive patch position themselves on the lensball. The spanned angle **θ_max_=170°.** Image (b) is a side view of the photoreceptor positions on the lensball. Image (c) shows a plan view of a photoreceptor distribution on the anterior hemisphere and image (d) in the posterior part of the retina. In a run of 66 simulations, the anterior photoreceptor distribution equals (μ=81.4; σ=5.9) versus the corresponding (μ=218.6; σ=5.9) in the posterior part of the retina.

In early evolutionary life forms, the photo sensitive organ can only register ambient light levels. Therefore, it does not matter whether the initial photoreceptor cell distribution in the “flat” light-sensitive region is uniform or randomly distributed. Figure 8 shows a computer simulation of how photoreceptor cells from a random 2D flat configuration distribute spatially across the lensball.

Both analyses (equation 1 and simulation) show that roughly 73% of the receptors are in the posterior part of the retina.

Due to this gooseneck photoreceptor density profile and the fact that the animal’s body shields the posterior photoreceptor cells from direct light activation, a dominant detection direction arises along the anterior-posterior axis of the lensball for incident light coming from the lateral hemisphere.

Light incident along this axis of the lensball will activate the posterior photoreceptor cells from within and - depending on the degree of refraction - create a blurred image of the outside world on the inside of the posterior half of the retina. The blurred image on the retina allows detecting the light-dark environmental pattern and lends directional sensitivity to the organ. The detectable spatial resolution in this pattern will depend on the regularity, the amount and spacing of the photoreceptors and the quality of the signal processing nerve center and forms the basis for a first primitive picture of the animal’s surroundings.

### The optical blur spot model

The refraction properties of a lensball and the photoreceptor distribution in the retina determine the maximum spatial resolution at which the outside world can be perceived. As the refractive index of the lensball starts to increase relative to the surrounding medium, it reduces the optical blur spot and the focal point becomes increasingly closer to the retina and thus improves the observable spatial resolution. To determine the structure of this blur spot as well as its maximum size, a mathematical model is developed in Box 3 (Figure 10).

The medium is initially water with a refractive index of 1.33, but can also be air with ~1.00 as a refractive index. The vitreous spherical lensball has an incipient refractive index of 1.34, which means that imaging in water is not yet a factor but in air it already leads to significant optical blur spot reduction.

In early evolutionary light-sensitive lensball, there are two effects that negatively affect unambiguous imaging.

The first effect is the activation of a photoreceptor in the posterior part of the retina. It can be direct or indirect producing a photoreceptor signal that is not unambiguously interpreted by the coordinating “brain” system regarding the direction of the light. This problem can be remedied completely by the evolutionary development of a lightproof outer layer on the posterior part of the organ.

Coverage of the lensball with a lightproof outer layer can occur up to at least halfway up the lensball (polar angle θ <90° in Figure 9) without limiting anteriorly incident photons from reaching the posterior retinal photoreceptor cells. The lightproof cover will now be called the “sclera”, while transparent part becomes the “cornea”. The light-sensitive spherical organ with a partially developed lightproof outer layer and a lensball is called a proto-eye.

**Figure 9.**
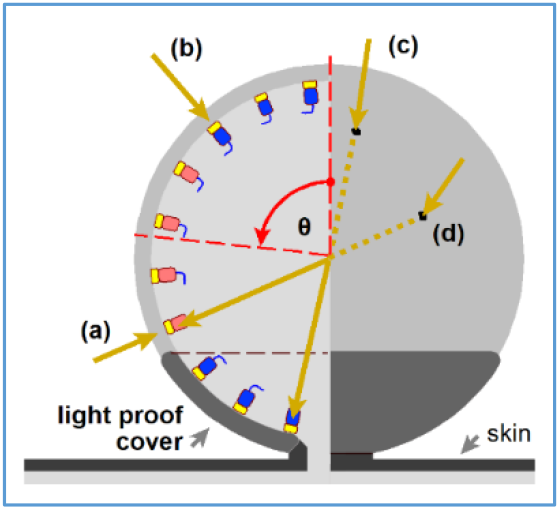
Diagram of a light-sensitive spherical organ with a partially developed lightproof outer layer and a transparent inner body. The yellow arrows represent how photons can reach a photoreceptor. The photoreceptor is symbolized by a rectangle with a light-sensitive tip (yellow). Photon (a) and photon (b) directly activate a photoreceptor (verted manner), whereas photon (c) and photon (d) enter the transparent vitreous body and via transmission through the vitreous body activate a photoreceptor in an inverted manner. The photoreceptor activated by photon (c) can only be activated in an inverted manner due to the presence of a lightproof shield. Photon (a) and photon (d) activate the same photoreceptor cell (red colored cell).

The second effect is the aberration on the posterior retina produced by refraction in the lensball, which causes the margins of anteriorly incident images to fold back over the central part of the image. This effect becomes more severe as the refractive indexes ratio n_2_/n_1_ increases and therefore is analyzed for aspects that may be an evolutionary driving force behind morphological changes improving the imaging properties of the light-sensitive organ. Equation 4 from Box 3 (Figure 10) is derived to calculate the contact point on the retina, for an incident photon, as a function of the angle of incident β, the polar angle θ and the refractive indices n_1_ and n_2_. The position of the impact point is expressed as the radius relative to the optical axis.

**Figure 10.**
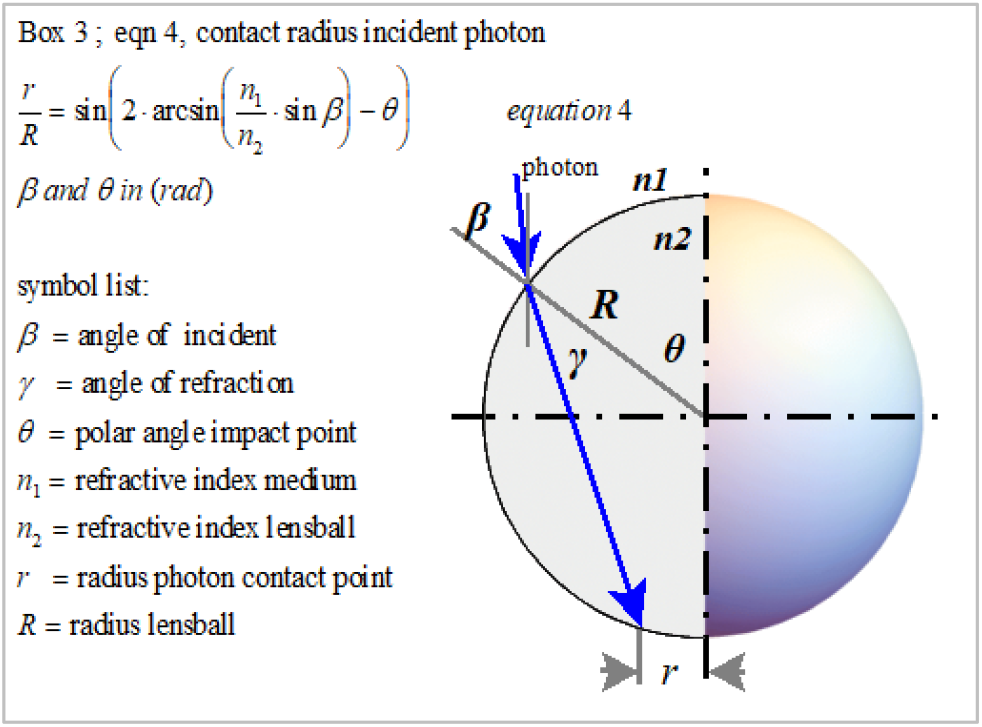
Box 3. Scheme for calculating the normalized retina contact radius r/R of an incident photon. Snell’s law: **n_1_** •sin(**β**)=**n_2_** •sin(**y**). For **β=θ**, the photon enters parallel to the polar axis. The polar angle **θ** of the impact point varies between 0 and π/2 radians. The photon contact point on the retina has a distance **r** to the optical lensball axis.

For the rest of this study, β=θ, meaning that photon incident is parallel to the optical axis. The normalized contact point radii are shown in Figure 11 as a function of the polar angle θ and the refractive indexes ratio n_2_/n_1_.

**Figure 11.**
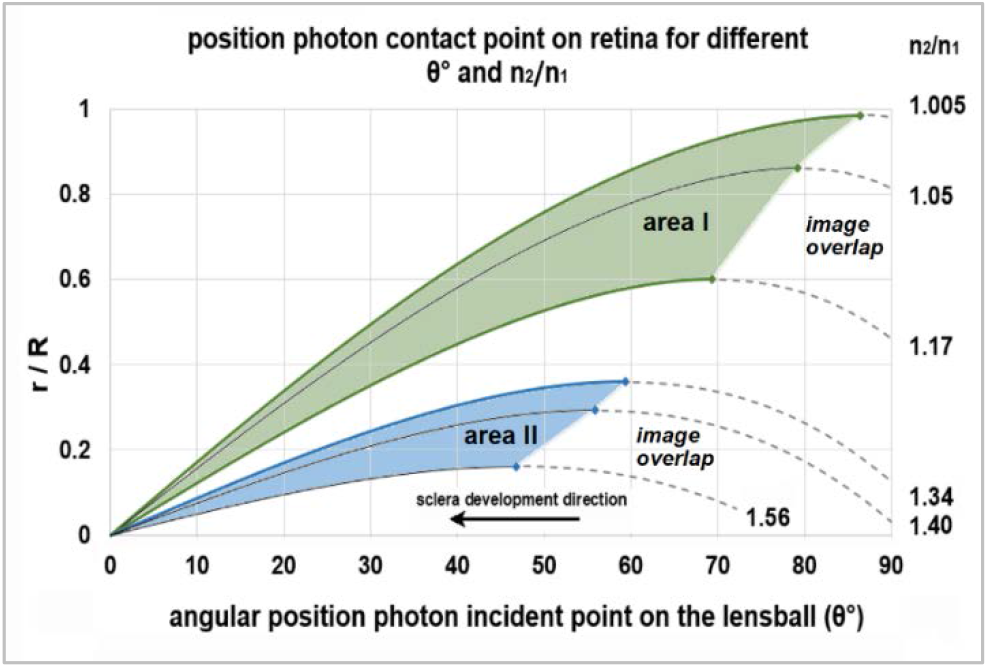
The normalized photon contact radius r/R as a function of the polar angle θ of photon incident point position and the ratio n_2_/n_1_. The dotted parts of the curves represent the incident angles that cause overlap in the blurred primitive image on the retina. Area I represents the achievable normalized optical blur spot radii of the proto-eye in water. Area II represents the situation in air. The higher the value of (n_2_/n_1_) and the smaller the aperture angle, the smaller the optical blur spot radius and the closer the focus point. The smaller the optical blur spot radius, the better the maximum spatial resolution.

With the given n_2_/n_1_ ratio and at increasing angles of incident *θ*, the photon impact point moves away from the optical axis until a maximum value is reached, after which it bends back toward the axis, creating a self-overlapping image. Blocking this overlap will provide a more readily interpretable image. This may have been the evolutionary trigger to further extend the sclera in the anterior direction to the aperture angle at which image overlap is eliminated. The rim of the sclera then forms the aperture for incident light. The aperture angle is defined as the angle between the line from the lensball center to the aperture edge and the optical axis. This aperture angle corresponds to the angle in Figure 11 that marks the maximum of a curve.

Depending on the ratio n_2_/n_1_, this corresponds to a sclera that can extend anteriorly to an aperture angle θ between 86° and 71° in water and in air between aperture angles corresponding to θ *=* 59° and 47°.

Further, the effect of the refractive index ratio on the normalized blur spot size was analyzed for various biological and environmental conditions (Figure 12). In water, where the index n_1_ = 1.33, the normalized optical blur spot radius (r/R) varies between 0.98 and 0.59, depending on the crystalline protein concentration, giving a n_2_ between 1.34 and 1.56 (Bassnett, Shi, & Vrensen, 2011). Figure 12 shows this process in the normalized maximum optical blur spot curve, where the maximum normalized optical blur spot radii from Figure 11 are plotted as a function of n_2_/n_1_.

**Figure 12.**
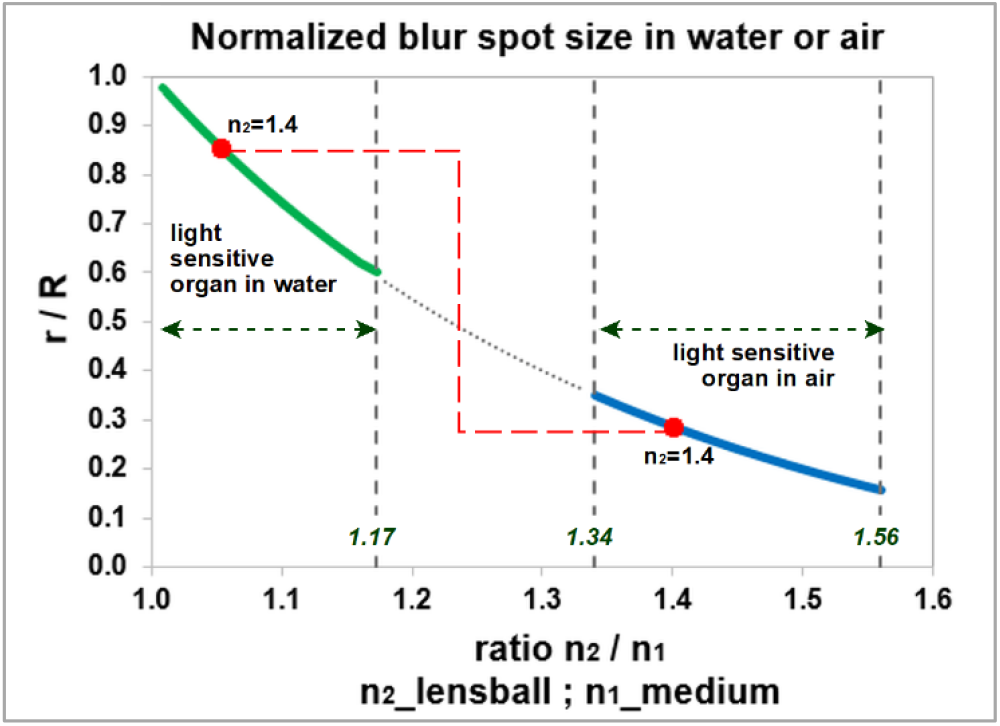
Normalized blur spot size as a function of the refractive index ratio n_2_/n_1_. The descending curve is determined by the maximum optical blur spot radii from Figure 7. The solid green curve applies to the underwater conditions (n_1_=1.33) and the solid blue curve to conditions in air (n_1_=1) under various values for n_2_ according to the crystalline concentration.

In air (n_1_=1), e.g. when the animal is foraging at or above the water surface, the normalized optical blur spot radius can shrink from 0.34 to 0.17 as the crystalline concentration is higher. For a refractive index n_2_=1.4, the optical blur spot radius decreases by a factor of 2.9 when the proto-eye crosses the water’s surface (shown as a red dashed line in Figure 12). The reduction in the blur spot radius averages a factor 3.2 +/- 0.4, suggesting that the fully developed inverted retina may have occurred first in water surface foraging animals.

The proto-eye with its cup-shaped inverted retina and the transparent lensball cap that functions as a positive lens allows the animal to perceive its surroundings with a coarse spatial resolution. The smaller the blur spot and the higher the photoreceptor density, the better the maximum detectable spatial resolution will become.

### The spatial resolution model

Improvement of detectable spatial resolution is recognized as an important trigger for eye evolution (Nilsson & Pelger, 1994). Their mathematical equation 5 (Figure 13. Box 4) can be used to determine the maximum detectable spatial frequency. The equation is designed for a proto-eye with an invaginated cup-shaped retina and a lens-free aperture for incident light, taking into account photon noise. Equation 5 is insensitive to verted or inverted photoreceptor activation. It assumes that the observable spatial resolution of the retinal photoreceptor mosaic is properly balanced with the optical quality of the rest of the light-sensitive organ.

**Figure 13.**
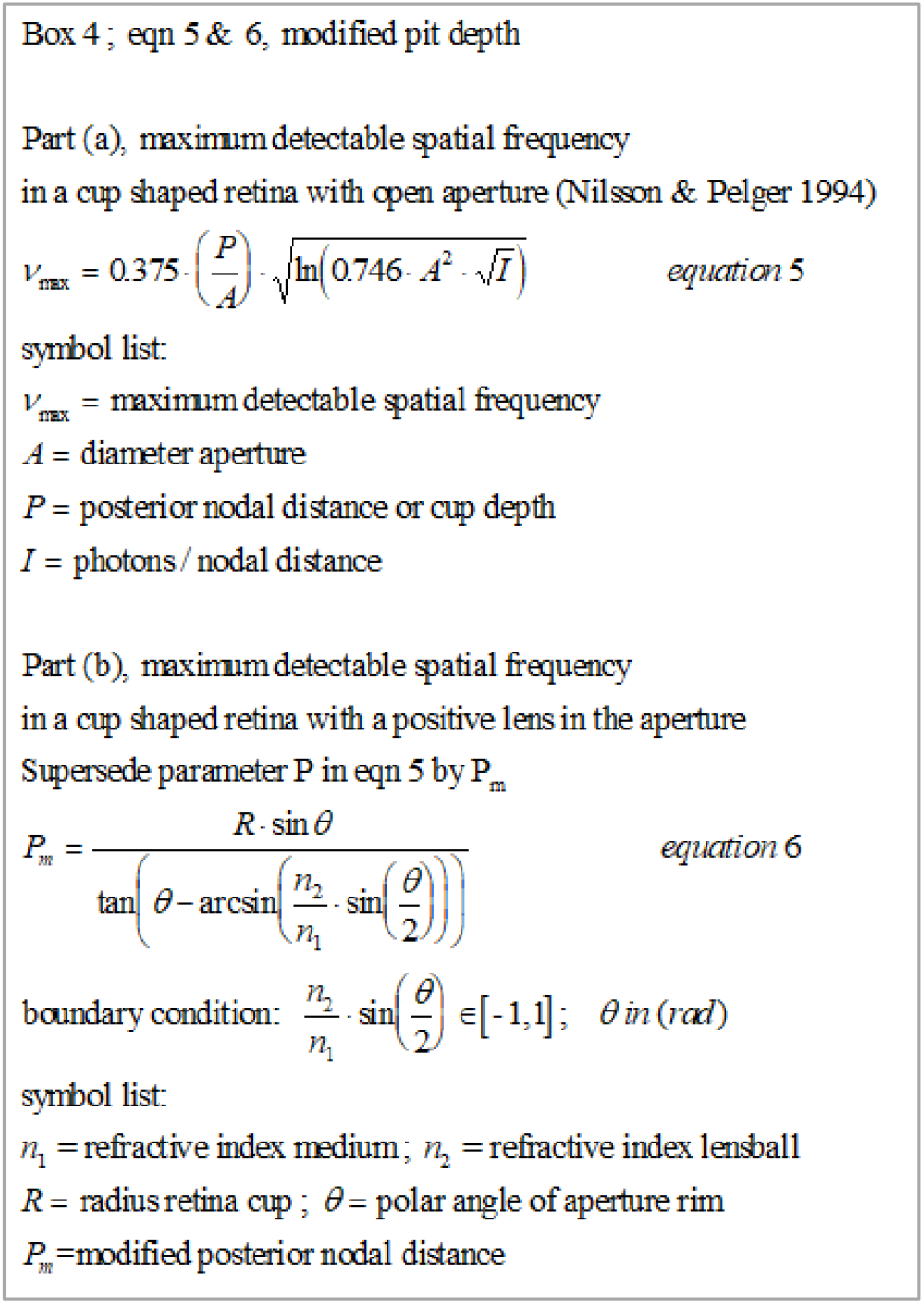
Box 4. Maximum detectable spatial frequency for a cup-shaped retina with a positive lens in the aperture. Part (a) presents the original equation taken from Nilsson and Pelger (1994). Part (b) presents the modified parameter **P_m_**, which accounts for the effect of the positive lens on the maximum detectable spatial frequency and replaces parameter **P** in equation 5. For the lensball, the relation between **P**, the cup radius **R** and polar angle **θ** of the aperture rim is **P**=**R**·(1+cos**θ**).

In equation 5, the maximum detectable spatial frequency is directly proportional to the posterior nodal distance (cup depth). This makes it possible to account for the positive lens effect in the aperture of the lensball by keeping the aperture constant and mathematically modifying the cup depth until the corresponding solid angle matches the solid angle for the center of the open-cup retina. The modified posterior nodal distance (modified cup depth) can be calculated with equation 6 (Figure 13. Box 4).

In Figure 14, the maximum resolution for the lensball is given as a function of the inverse of the normalized cup depth Pm/A for the variable refractive ratio n_2_/n_1_, following Nilsson and Pelger (1994). In water, this provides an increase of ~20% in maximal achievable spatial resolution at an aperture angle of 18°. At the aperture angle of 18°, this yields a ~46% increase in spatial resolution in air. The spatial resolution can be further increased by reducing the aperture for incident light, i.e. through expansion of the sclera and an increase of the refractive index n_2_ untill a maximum is reached for an aperture angle of 11°.

**Figure 14.**
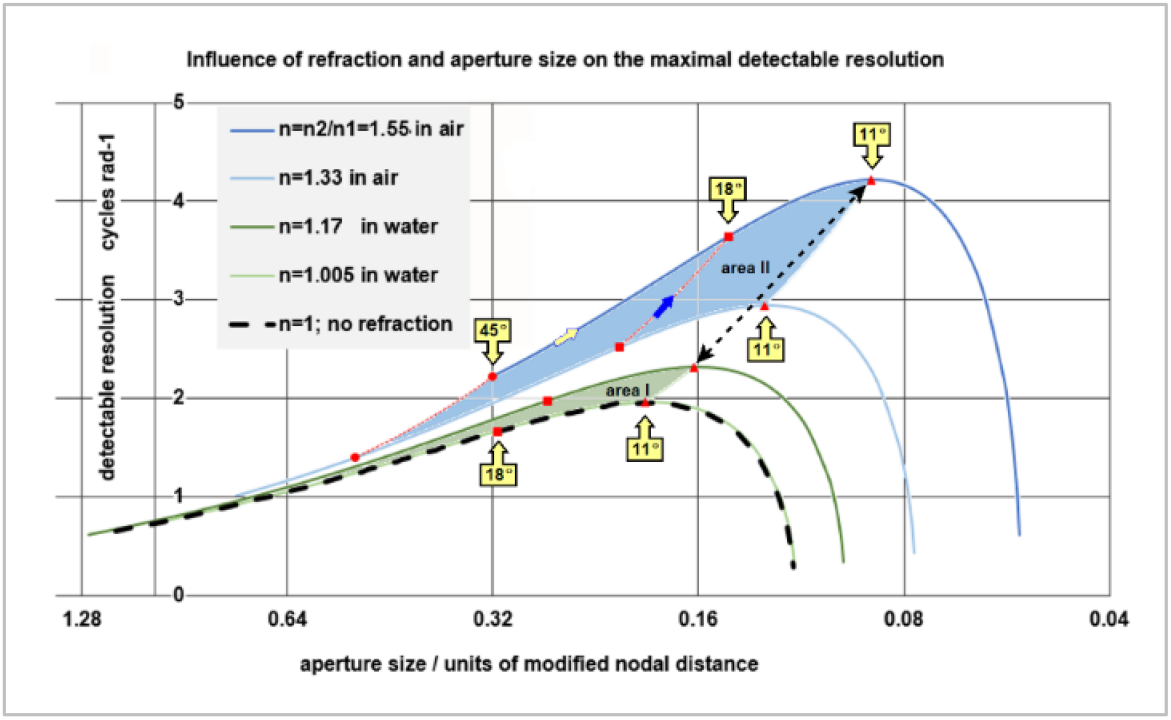
Options for improvement of detectable resolution for the scenario A=Ac and θ_max_=170°. The influence of the refractive index of the lensball is calculated for a light intensity of 10^4^ photons per normalized surface per second per steradian, taken from Nilsson and Pelger (1994). The solid lines enclose the area within which the evolutionary optimization of the proto-eye in respect to the spatial resolution can take place. Area I represents the achievable spatial resolution in water. Area II represents the situation in air. The dashed bottom line represents the achievable spatial resolution for the open-cup proto-eye and the lensball proto-eye, provided that both eye types have the same aperture size and the refractive indices ratio **n_2_**/**n_1_** equals one^2^. The red markers on the lines illustrate progress in detectable resolution as a consequence of aperture angle reduction. The blue arrow points in the direction of an increase in detectable spatial resolution by enhancing the refractive index of the lensball. The bi-directional black arrow represents the jump in spatial resolution when the proto-eye emerges above the water.

A further increase in maximum detectable spatial frequency is possible by developing extra eye appendages that increase the amount of light entering the aperture to reach the retina. This can be achieved by a positive lens in the aperture or the lens like outer cornea.

### Eye appendages

For the sake of convenience, the proto-eye on a stalk is referred to as a “stalk eye” from here onwards. The stalk eye has refractive properties on its spherical surface that allow it to focus, but in air and with a vitreous body with a refractive index of 1.56, the focal point is still behind the retina, resulting in blurred imaging. For sharp imaging, the focal point must be on the retina, which will require the development of additional appendages such as a cornea and/or lens.

To further improve spatial resolution, the proto-eye must evolve appendages such as a lens and cornea. The formation of these appendages should not conflict with the configuration of the proto-eye and should be able to develop without significant morphological change. The main eye appendages are discussed in Figure 15.

**Figure 15.**
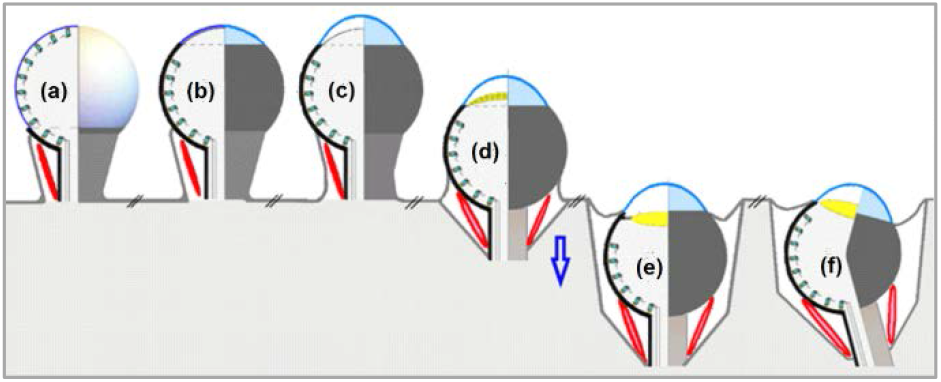
Color-coded hypothetical eye appendages and their characteristics. a) The stalk containing a channel for nerves and veins, with external ocular muscles (shown in red) b) Translucent anterior aperture, free of photoreceptors, coupled to an extended photoreceptors, coupled to an extended lightproof sclera forming a proto-eye c) Protruding cornea (the proto-eye gets features of an eye) d) Proto-lens and retreating apparatus.(symbolized by the blue arrow) e) Retreated eye with lens f) Tilted optical axis positioning the blur spot on the retina

**Extra ocular muscles** (Figure 15a-f). The eye stalk places the visual organ farther away from the body and if the stalk is equipped with extra-ocular muscles, the muscles allow the eye to be oriented. Both features contribute to achieve a wider field of view. The stalk of the proto-eye contains an optical channel.
**Frontal eye aperture and extension of the sclera** (Figure 15b). The narrow translucent opening bordered by an extended sclera allows rays of light to penetrate in such an angle through a vitreous body that an univocal detailed image can be produced in the dense light-sensitive layer in the posterior part of the eye.
**Cornea** (Figure 15c). The few photoreceptors in the transparent anterior part of the retina become inferior to the photoreceptors in the posterior retina, likely prompting their disappearance as evolution progresses. The transparent epithelial layer covering the anterior retina layer extends from the sclera and by bulging forming a chamber filled with an aqueous humor. This transparent epithelial layer or cornea has a stronger curvature than the lensball and provides an additional refractive surface in air that reduces the blur spot even further, which contributes to better imaging. However, a cornea does not contribute to improved vision in water. The difference in the refractive index of water and the aqueous humor is too small. When the cornea and sclera become rigid structures, they provide mechanical support to the eyeball. Another consequence of the stiffness of the cornea is that its normalized refractive power is a fixed value. In water, a lens is required to realize a more focused projection on the retina.

It is supposed in this article that tissue areas with different biological functions are separated by basal membranes. Basal membranes are very old morphological structures from an evolutionary perspective. In the embryonic phase, they can fuse and separate, creating new cell regions with different biological functions (Kefalides & Borel, 2005; Morrissey & Sherwood, 2015). These characteristics of basal membranes play a role in lens development.

**Lens** (Figure 15 d/e). The photoreceptors in the anterior part of the proto-eye disappear, leaving a transparent epithelial cell layer between two basal membranes. This cell layer can thicken to gain a lenticular shape. Consequently, the incoming rays of light converge to focus on the retina, whereby the observable spatial resolution is optimized. This type of primitive lens occurs in parietal eyes. The basal membranes constrict and fuse, which can lead to a separation of the lens from the retina but they are anchored to each other by fibrils. A lens has the advantage of a changing curvature and changing refractive index, causing the lensball to lose its function as a refractive surface for incident light. Maintaining a high concentration of crystalline proteins in the vitreous lensball is no longer necessary.
**Retraction of the proto-eye into the body.** The protruding soft-tissue stalk eye in the predecessor of the vertebrates is vulnerable and therefore probably retracted into the body, while the eyeball tilt and rotation and field of vision are retained.

Note that a structure that generates an additional shrinkage of the aperture for the incident light - like an iris does - would be beneficial for the visual resolving power, but it would not be necessary for the functioning of the proto-eye.

## Results

### The photoreceptor density model

The developed local photoreceptor density equation predicts a low density anterior and a high density posterior, allowing the posterior retina to develop in an inverted retina sensitive to incident anterior light, creating the basis for detectable spatial resolution.

### The optical blur spot model

The model offers two triggers for sclera development and a trigger for aperture clearance of light-sensitive cells.

Prevention of photoreceptor signals that the coordinating “brain” system cannot unambiguously interpreted, Figure 9 by coverage of the posterior part of the lensball with a lightproof outer layer from stalk (θ~170°) to an aperture with a polar angle θ ≤ 90°.

Annihilating the negative effects of aberration can be achieved by extending the sclera further in the apex direction up to a polar angle that corresponds with the maximum blur spot radius depending on the environment air (59° ≤ θ ≤ 47°) or water (86°≤ θ ≤ 71°).

### The spatial resolution model

The spatial frequency model of Nilsson and Pelger (1994) for an open-cup retina can be used for a lensball cup retina with a positive lens surface in the aperture by modifying the parameter pit depth. An equation for calculating this modified pit depth is presented in Box 4. The spatial frequency model allows further extending the sclera in apex direction up to a polar angle θ =11°, which corresponds with the maximum detectable spatial frequencies in Figure 14. This angle is independent of the air or water environment and the refractive index of the lensball.

**Figure 16.**
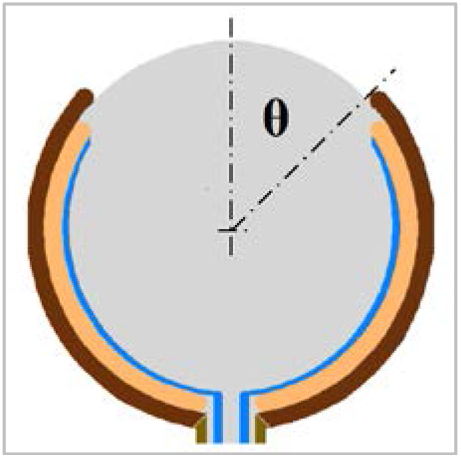
Color-coded schematic overview of the proto-eye. The grey area represents the humorous lensball, the blue lines the nerves towards the coordinating “brain” system, the ochre band the retina and the brown band the light-tight sclera. Polar angle **θ** determines the aperture.

The proto-eye with the full developed sclera and corresponding aperture has a detectable spatial resolution in air between three and four cycles/rad and in water around two cycles/rad, proving that the evagination scenario for eye development has the full potential to explain the origin of the inverted retina.

Figure 17 presents the hypothetical developmental pathway for the inverted retina and lens-eye appendages of the vertebrate proto-eye, parallel to an accepted developmental pathway of the mollusc proto-lens-eye. It shows the embryological formation of the vertebrate eye and the involvement of the tool kit genes Pax6 and Pax2.

**Figure 17.**
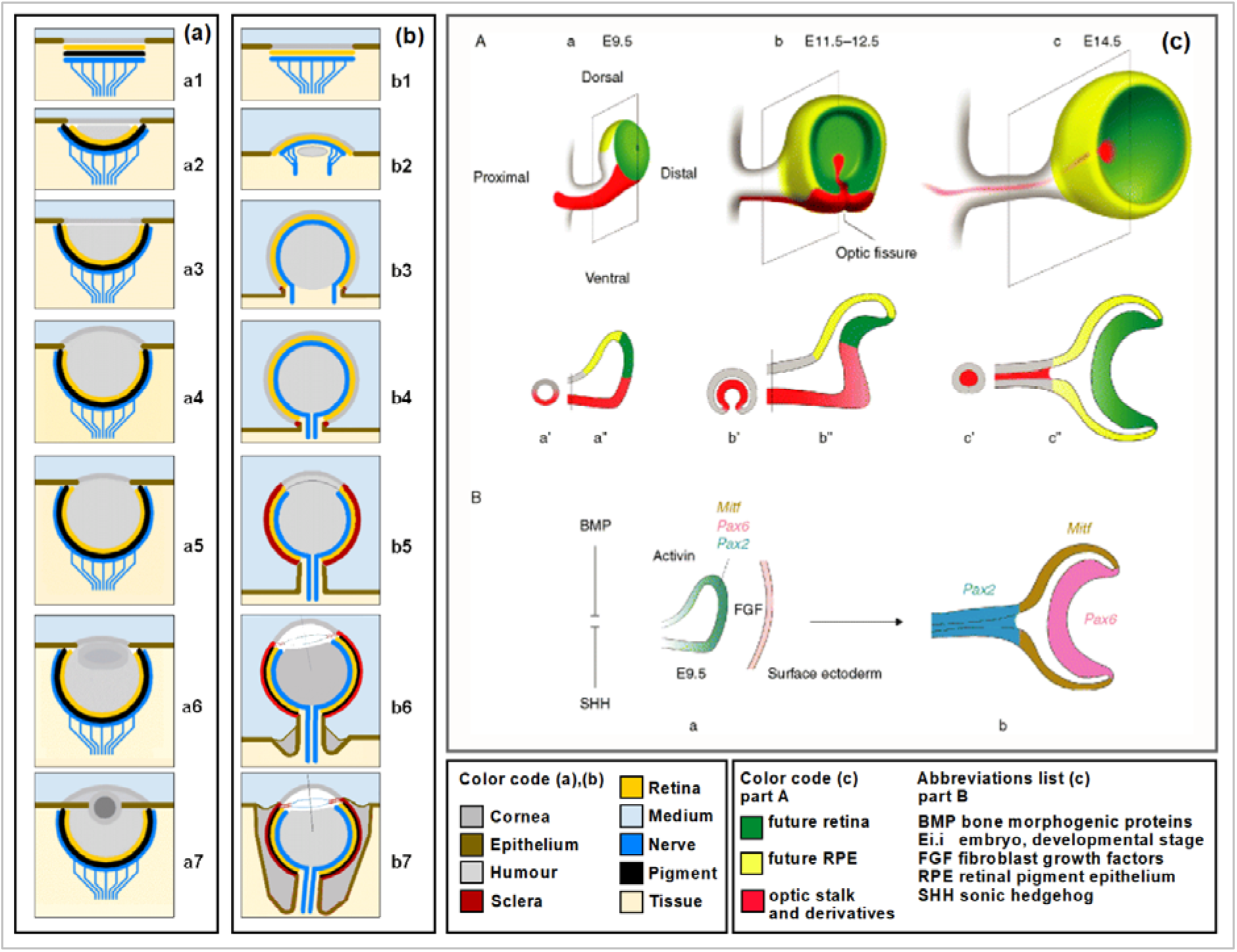
Schematic comparison of the hypothetical evolutionary pathways of two different proto-eyes. Figure 17a, steps 1-7 of the developmental pathway of the mollusc proto-lens-eye, redrawn from Nilsson and Pelger (1994). Figure 17b, developmental steps of the vertebrate predecessor proto-eye. b1, The initial light-sensitive patch has a retina with primitive photoreceptors and not yet a screening pigment layer compared to (a1) in the evolutionary sequence given by Nilsson and Pelger (1994) and Land and Nilsson (2012). b2, Different evolutionary pathways for evaginating photoreceptor patch compared to the invaginating patch a2. b3, Development of posterior retina. b4, Posterior inverted retinal activation, start detectable spatial resolution and imaging. b5, Fully developed inverted retina, light-tight sclera and clear aperture shaping the vertebrate proto-eye. b6. Speculative; Primitive lens and screening pigment layer development. The optical axis tilts with respect to the stalk axis. Starting retraction of the proto-eye into the body. b7, Speculative; Fully developed proto-eye in the predecessor of the vertebrates. Figure 17c, visualization the areas where the tool kit genes Pax2 and Pax6 come into action in the human embryo. It illustrates the close involvement of the Pax2 gene in optical nerve tube development but also in blind spot development (red area) and the involvement of Pax6 gene (green area) in the retina cup development in the vertebrate eye, original caption of the figure. Reprinted from “The other pigment cell: specification and development of the pigmented epithelium of the vertebrate eye,” by K. Bharti, M. T. Nguyen, S. Skuntz, S. Bertuzzi, and H. Arnheiter, 2006, Pigment cell research, 19(5), p. 382. Copyright 2006 by John Wiley and Sons. Reprinted with permission.

## Discussion

It is likely that an inverted retina may have evolved in the predecessors of the vertebrates from a primitive spherical light-sensitive organ on a stalk. The transparent spherical light-sensitive organ has a retina that is verted activated by direct light and inverted activated by incident light. Light-sensitive organs on a stalk are a common feature in nature. This study shows that small configurational modifications triggered by spatial resolution optimization give rise to an inverted retina in a functional proto-eye by using incident light for all vision functions rather than direct light. Incident light captured by the concave posterior part of the retina allows for imaging, whereby the retina in the posterior part becomes superior to the anterior part. Configuration (b5) from Figure 17 represents this proto-eye.

The optimum aperture is independent of the ratio n_2_/n_1_ and has a polar angle of 11°. This suggests that for the stalk eye, the sclera development has priority over other evolutionary options for spatial resolution improvement. The maximum resolvable spatial frequency that can be achieved with the optimum aperture varies in water between 2 and 2.3 (cycles per radian) and in air between 3 and 4.3 (cycles per radian), depending on the protein concentration and therefore the refractive index n_2_ of the vitreous body. A deep-sea isopod has a maximum resolvable spatial frequency of 1.9, and it is 3.6 for a Nautilus (Land & Nilsson, 2012).

Another reason is that in the spherical proto-eye, photoreceptor density across the sphere increases by a factor of ~16 from anterior to posterior. This factor is independent of the chosen invagination scenario but based on the assumption that the photoreceptor cells retain their position in the support matrix during invagination.

The assumption that the photoreceptor cells retain their position in the support matrix upon bulging of the light-sensitive organ and therefore become unevenly distributed is vulnerable, because there may be an early evolutionary genetic mechanism that keeps the photoreceptor cells evenly distributed across the surface area at all times during the shape change process. This mechanism is not expected to occur in an organ sensitive only to light intensity and light direction and in which detectable spatial resolution does not yet play a role. However, it is to be expected in a proto-eye that already has imaging capabilities, as a more uniform retinal photoreceptor pattern directly affects image resolution positively.

Photoreceptors located in the convex anterior part cannot contribute to detectable spatial resolution because the spatial angle of view is close to 180**°** and therefore the detectable spatial resolution is zero. The processing of the fewer signals coming from the anterior part of the retina loses its function and the corresponding light-sensitive cells will disappear evolutionarily by genetic drift. According to Sallan, Friedman, Sansom, Bird, & Sansom (2018), this usually requires few generations in isolated populations. Tidal zones and coastal areas are prime areas for small isolated populations. The remains of water bound plants and animals wash up on the coast and the banks of rivers and waterholes. It is expected that these zones were already very bioactive in the pre-Cambium period.

An increase of photoreceptor density in the posterior part of the retina and more regular distribution of the photoreceptors will further increasing spatial resolution. In this regard, a hexagonal configuration is the most obvious, because it yields the densest photoreceptor bunching and thus the smallest spacing between photoreceptors, as we can observe in contemporary animal eye retinas.

The soft-tissue stalk eye is vulnerable to physical damage and therefore probably retracted into the soft-tissue animal body while still being able to scan the animal’s surroundings. During the Cambrian, hard internal and external body parts develop in a variety of animal species and embed the vertebrate eye into the skull. The eye and optic stalk are even better protected,

There are no known fossil finds from the Precambrian and Cambrian that can support the stalk eye hypothesis in vertebrates. It is possible that the notion that stalk eyes could be the precursor to the vertebrate eye contributes to the interpretation of eye characteristics in pre-Cambrian fossils. Evidence of the soft-tissue proto-eye on a stalk can possibly be found in chemical traces of retinal molecules beyond the body imprint.

The amphioxus eye - with its inverted interpreted retina - does not fit directly into the stalk eye hypothesis, unlike the more advanced, bulging eyes of Conodonts (Donoghue, Forey, & Aldridge, 2000). The eye of the Hagfish may represent phase 3 (Figure 2) because it comprises an inverted retina around a vitreous spherical body but has no screening pigment layer (Dong & Allison, 2021). Next, the eye was retracted into the body without maintaining the optic stalk. The eyes in the larval phase of the Lamprey are similar to the eye of the Hagfish but migrate outward with the development of the lens and cornea.

The stalk is a new structure requiring a genetic basis that may be provided by the Pax2 gene, which plays an important and tightly delineated role in the embryonic development of the optic tract in the vertebrate eye (Torres, Gómez-Pardo, & Gruss, 1996). In his review of the various genes involved in eye development, Fernald (2004) indicates that the Pax2 gene is expressed within the embryonic vertebrate eye only and not in other phyla. This is in contrast to the Pax6 gene, which is active in all phyla. Fernald (2004) postulates that Pax6 was almost certainly active in the precursors of vertebrates, shellfish, insects and molluscs, while it is unclear whether the same is true for the Pax2 gene. The Pax2 gene belongs to the early evolutionary genetic toolbox, along with - among others - the Pax6 gene, which makes it likely that Pax2 was involved in the development of the proto-eye on a stalk at an early phase of evolution. No direct conflict between embryonic development and morphological outcome can be found. The initial retina in the light sensitive patch is of ectodermal origin and therefore the retina in the proto-eye. The ectodermal retina probably transformed later into neuroectodermal tissue.

There are extant fish species that develop full stalk eyes in their larval phase (Weihs & Moser, 1981). These types of larval eyes can potentially elucidate the genetic process and explain which genes are responsible for the larval eye stalk. If involvement of the Pax2 gene is demonstrated, this would serve as a strong indication that this developmental pathway may also have occurred at an early evolutionary phase and was genetically conserved. Otherwise these fish stalk eyes are a more recent evolutionary development.

From a topological point of view, there is a need to rethink the roles of Pax2 and Pax6 in vertebrate retina development. The involvement of Pax2 by indenting the budding optic vesicle ventrally results in the development of the optic stalk and the vertebrate retina with the optic disk (Figure 17c. Bharti et all., 2006). This means that topologically the future retina region on the optic vesicle invaginates as an incomplete spherical cup with a wedge-shaped opening that deforms to a spherical segment during closing of the retinal fissure. With that the development of the retina in the vertebrate eye is the combined result of two different topological actions; ventral indentation and distal indentation. One could speculate that ventral indentation only leads to the development of a functioning inverted retina in a proto-eye like the eye of the hagfish. Distal indentation only will not lead to development of the optic disk - blind spot - and therefore not to a functioning inverted retina.

An intriguing question is whether the present mammalian eye with its optic stalk and extrao-cular muscles is a further developed proto-eye on a stalk that - due to its fragility - was gradually retracted into the animal’s head in its entirety and while retaining its directional power due to evolution. In this case, the contemporary optic stalk in the mammalian eye could be a functional successor of the proto-eye stalk.

## Conclusion

An evaginating flat light-sensitive area with spaced-out photoreceptor cells in a “verted” - light facing - orientation can develop into a outwardly protruding spherical transparent light-sensitive organ covered with a light facing retina.

Whether the retina in the outwardly protruding spherical transparent light-sensitive organ should be considered “verted” or “inverted” depends on the direction from which light activates the light-sensitive retinal cells. The transparent spherical light-sensitive organ intrinsically allows for both, as is shown in Figure 9.

Blocking photoreceptor activation by direct light and accepting incident light to reach and activate them restricts the retina to the inverted activation mode without any need for morphological retina modifications.

This thus makes it a light-sensitive organ from which the “inverted retina” in vertebrates may have arisen under evolutionary pressure of achieving the highest possible detectable spatial resolution.

Assuming that the inverted retina developed before neurulation occurred (Carreras, 2018), this study suggests that the primitive retina in the predecessor of vertebrates was of ectodermal origin and neuroectodermal after neurulation.

This study also shows that the retina probably developed along two scenarios, namely an invagination process resulting in a verted retina for invertebrates and an evagination process resulting in the inverted retina for vertebrates, all before a proto-eye came into existence.

It also shows that in case of lacking fossil evidence, insufficient intermediate species or not-yet-available genetic evidence, physical modeling has its use as a tool to analyze the possible origin of the inverted retina in vertebrates.

## Acknowledgments

I would like to extend my thanks to José M. de Veer, Dr. Stef M. Olsthoorn, Drs. Fred A.J.M. Neelissen, Dr. Petra van Langevelde and Dr. E.S. Pierson for helpful discussions and their valuable comments on the manuscript.

## Footnotes

1 *Due to the mirror symmetry of the evagination and invagination process, equation 1 can be applied to both sphere geometry and cup geometry*.

2 *Spatial resolution values calculated with equation 5 of Nilsson and Pelger (1994) are two times higher than the values presented in Figure 1b in their article. Despite this discrepancy, it is assumed that their equation 1 can be used to analyze spatial resolution for an open cup-shaped retina. The layout of Figure 14 is taken the same as the layout of their Figure 1b*.

## References

1. Bharti, K., Nguyen, M. T. T., Skuntz, S., Bertuzzi, S., & Arnheiter, H. (2006). The other pigment cell: specification and development of the pigmented epithelium of the vertebrate eye. Pigment cell research, 19(5), 380–394.

2. Bassnett, S., Shi, Y., & Vrensen, G. F. (2011). Biological glass: structural determinants of eye lens transparency. Philosophical Transactions of the Royal Society B: Biological Sciences, 366(1568), 1250–1264.

3. Carreras, F. J. (2018). The inverted retina and the evolution of vertebrates: an evo-devo perspective.

4. Dawkins, R. (1997). Climbing mount improbable. WW Norton & Company.

5. Dong, E. M., & Allison, W. T. (2021). Vertebrate features revealed in the rudimentary eye of the Pacific hagfish (Eptatretus stoutii). Proceedings of the Royal Society B, 288(1942), 20202187.

6. Donoghue, P. C., Forey, P. L., & Aldridge, R. J. (2000). Conodont affinity and chordate phylogeny. Biological Reviews, 75(2), 191–251.

7. Fernald, R. D. (2004). Evolving eyes. International Journal of Developmental Biology, 48(8-9), 701–705.

8. Kefalides, N., & Borel, J. (2005). Morphology and ultrastructure of basement membranes. Curr. Top. Membr, 56, 19–42.

9. Land, M. F., & Nilsson, D. E. (2012). Animal eyes. OUP Oxford.

10. Morrissey, M. A., & Sherwood, D. R. (2015). An active role for basement membrane assembly and modification in tissue sculpting. Journal of cell science, 128(9), 1661–1668.

11. Nilsson, D. E. (2009). The evolution of eyes and visually guided behaviour. Philosophical Transactions of the Royal Society B: Biological Sciences, 364(1531), 2833–2847.

12. Nilsson, D. E., & Pelger, S. (1994). A pessimistic estimate of the time required for an eye to evolve. Proceedings of the Royal Society of London. Series B: Biological Sciences, 256(1345), 53–58.

13. Sallan, L., Friedman, M., Sansom, R. S., Bird, C. M., & Sansom, I. J. (2018). The nearshore cradle of early vertebrate diversification. Science, 362(6413), 460–464.

14. Torres, M., Gómez-Pardo, E., & Gruss, P. (1996). Pax2 contributes to inner ear patterning and optic nerve trajectory. Development, 122(11), 3381–3391.

15. Weihs, D., & Moser, H. G. (1981). Stalked eyes as an adaptation towards more efficient foraging in marine fish larvae. Bulletin of Marine Science, 31(1), 31–36.

